# An Agentic Platform for Drug Repurposing Unified across Molecular, Phenotypic, and Clinical Scales

**DOI:** 10.64898/2026.04.19.719462

**Authors:** Cheng Wang, Mohamed El Moussaoui, Dongdong Zhang, Prathiksha Prabhakaraalva, Serge Merzliakov, Rita Jui-Hsien Lu, Nabila Zaman, Goutam Chakraborty, Kuan-lin Huang

**Author notes:** To whom correspondence should be addressed. Kuan-lin Huang.

## Abstract

Drug repurposing offers an effective path to new therapies, yet existing computational approaches rely on a single line of evidence and are rarely validated across biological scales. We present LinkD, an integrated framework that unifies diffusion-based affinity prediction, proteome-wide selectivity scoring, phenotypic validation, and population-scale clinical evidence. LinkD-Bind predicts binding across 14,981 drugs and 20,385 human targets, ranking first in 8 of 9 BindingDB, Davis, and KIBA evaluations, with the largest gains under cold-start conditions. LinkD-Select recovers 95.3% of known drug-target pairs by combining selectivity scoring and molecular docking. LinkD-Pheno integrates drug-sensitivity and CRISPR dependency data across 960 cancer cell lines, identifying 34 novel drug-gene pairs and recovering ∼85% of known targets among the top 50 candidates. Across 11.5 million individuals from Mount Sinai and UK Biobank, LinkD-prioritized β-blockers propranolol (HR 0.82) and carvedilol (HR 0.92) reduced 5-year prostate cancer incidence relative to metoprolol, corroborated by ADRB2 docking and LNCaP growth inhibition. LinkD-Agent, which can effectively orchestrate all evidence layers, is served on a publicly available web platform (https://linkd-agent.onrender.com/), enabling a wide range of users to derive new drug repurposing opportunities through natural language queries.

## Introduction

The identification of new therapeutic indications for approved drugs, known as drug repurposing or repositioning, has emerged as one of the most efficient strategies for expanding the pharmacological arsenal against complex diseases^1^. Because repurposed compounds have established safety and pharmacokinetic profiles, they can enter clinical trials more rapidly and at substantially lower cost than *de novo* candidates, shortening timelines from discovery to patient benefit by years^2^. Several landmark examples, including the repositioning of thalidomide for multiple myeloma and sildenafil for pulmonary hypertension, underscore the clinical and economic value of this paradigm^3^. Despite these successes, systematic repurposing at scale remains hindered by fundamental methodological limitations.

Current computational repurposing strategies can be broadly grouped into target-centric and phenotype-driven approaches, each with distinct strengths and blind spots. Target-centric methods predict drug-target interactions based on molecular structure, docking simulations, or sequence-level features, and have benefited substantially from advances in deep learning^4–6^. Deep learning models such as DeepDTA^7^, GraphDTA^8^, and DeepPurpose^9^ learn joint representations of ligand and protein inputs to predict binding affinity with increasing accuracy. However, these methods typically evaluate interactions in isolation, assigning a single affinity score to each drug-target pair without accounting for the broader selectivity landscape, the distribution of a compound’s affinities across the proteome, which is critical for anticipating both therapeutic efficacy and off-target liabilities^10^. Existing metrics, including entropy-based scores and Gini coefficients,^11,12^ have only been applied primarily to kinase inhibitor panels and are rarely incorporated into affinity prediction pipelines as a primary modeling objective. Conversely, phenotype-driven approaches leverage large-scale perturbation datasets, including pharmacological sensitivity screens and CRISPR-based genetic dependency maps, to infer functional relationships between drugs and cellular states^13–15^. While these methods capture disease-relevant biology that structure-based models may miss, they lack the mechanistic resolution needed to pinpoint specific molecular targets and are constrained by the coverage of available screening datasets^16^.

A further gap exists in the translation of molecular and cellular predictions to clinical relevance. The vast majority of repurposing studies terminate at the computational or *in vitro* stage, with few systematically evaluating whether predicted interactions manifest as detectable clinical signals in patient populations^17^. The growing availability of population-scale electronic health record (EHR) systems and biobanks now offers an unprecedented opportunity to assess drug-disease associations at the epidemiological level^18,19^. Integrating such real-world evidence with molecular predictions could substantially strengthen the evidentiary basis for repurposing hypotheses, yet few existing frameworks attempt this integration in a systematic manner.

Here we introduce LinkD, a unified computational framework that addresses this gap through four tightly integrated modules. LinkD-Bind learns a shared latent representation of drugs and targets from molecular structures and protein sequences, then refines this representation through a denoising diffusion process to model interaction uncertainty and generalize to previously unseen compounds or proteins. LinkD-Select evaluates predicted affinities across the full proteome, quantifying selectivity using entropy-based and complementary distributional metrics that explicitly distinguish on-target preference from off-target dispersion. LinkD-Pheno contextualizes and validates these selectivity predictions through concordance analysis of large-scale pharmacological sensitivity profiles and CRISPR gene-dependency data, linking molecular interaction predictions to functional cellular consequences across tissue lineages. LinkD-Agent implements the framework as an interactive AI agent that enables transparent, evidence-based, and query-driven aggregation of mechanistic, phenotypic, and clinical evidence, including population-scale EHR validation across 11.5 million individuals, for scalable hypothesis generation and drug repurposing.

## Results

### The LinkD Framework that Combines Target-Centric and Phenotypic Approaches

LinkD is an integrated framework that uniquely cobmines target-centric and phenotypic drug discovery through four modules: diffusion-based affinity prediction (**LinkD-Bind**), proteome-wide selectivity scoring (**LinkD-Select**), phenotypic concordance analysis (**LinkD-Pheno**), and interactive multi-evidence integration (**LinkD-Agent**) (**Fig. 1a**; Methods). For a given drug-protein pair, LinkD-Bind encodes molecular structure and protein sequence into a shared latent space, refines the joint representation through a denoising diffusion process, and predicts binding affinity across a pan-proteome panel of 20,385 human protein targets. This proteome-wide affinity landscape provides the foundation for downstream selectivity analysis, phenotypic validation, and clinical corroboration.

**Figure 1.**
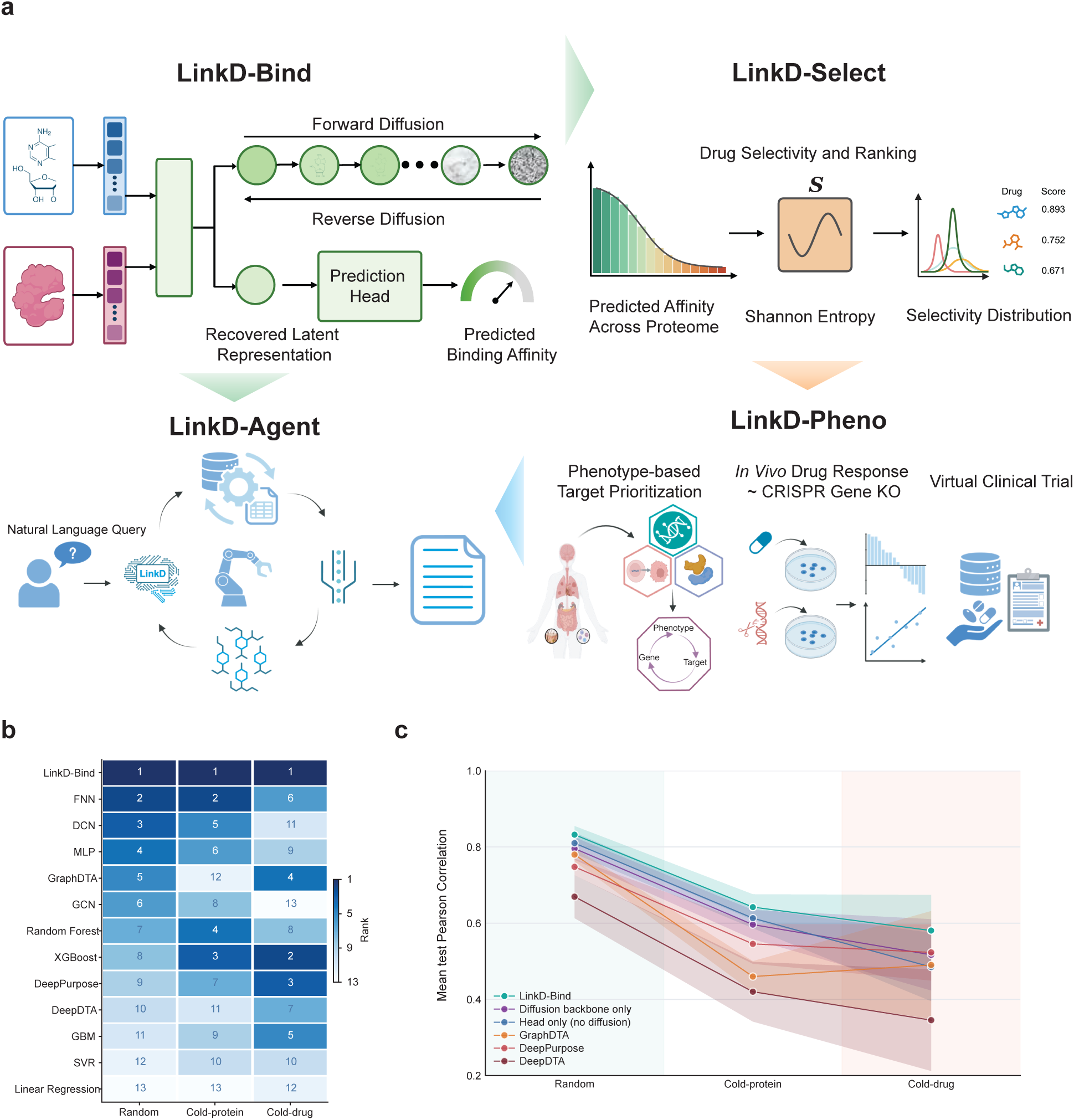
The LinkD framework integrates proteome-wide affinity prediction, selectivity scoring, phenotypic validation, and clinical evidence for drug repurposing, with LinkD-Bind outperforming established drug-target affinity predictors on standard benchmarks. **a.** Overview of the four LinkD modules. LinkD-Bind encodes small-molecule drugs from SMILES strings using ChemBERTa and protein targets from amino-acid sequences using the ESM2 protein language model, projects both into a shared latent space, and refines the joint representation through a forward and reverse denoising diffusion process that models interaction uncertainty. A hybrid MLP and Random Forest prediction head then estimates pairwise binding affinity (pKd) for 14,981 drugs across 20,385 human protein targets. LinkD-Select computes proteome-wide selectivity from the predicted affinity distribution of each drug. An entropy-based score, together with complementary concentration and affinity-gap metrics, distinguishes compounds with focused on-target binding from those with dispersed off-target profiles. Structure-level validation through binding pocket identification and molecular docking provides orthogonal support for predicted selectivity. LinkD-Pheno contextualizes selectivity predictions using pharmacological sensitivity profiles and CRISPR-Cas9 gene-dependency scores across 960 cancer cell lines spanning 13 cancer tissue lineages, and extends this validation to population-scale real-world evidence from electronic health records in 11.5 million individuals across the Mount Sinai Health System and UK Biobank cohorts. LinkD-Agent exposes all modules through an interactive interface that decomposes natural-language queries into structured, auditable analytical steps. b. Rank-based benchmarking of LinkD-Bind against twelve baseline drug-target affinity predictors averaged over the BindingDB, Davis, and KIBA benchmark datasets. Each cell reports the rank of one model in one split (1 = best, 13 = worst); darker shading indicates higher rank. LinkD-Bind ranks first in all three splits and in 8 of 9 (dataset × split) combinations overall. c. Mean test-set Pearson correlation across the three benchmark datasets for LinkD-Bind, two ablated variants (diffusion backbone only; prediction head only, no diffusion), and three deep-learning baselines, under each split. Shaded bands indicate the dataset-to-dataset variability. Both ablated variants underperform full LinkD-Bind across all splits, with the largest gaps appearing under cold-protein and cold-drug conditions, demonstrating that diffusion-based latent refinement and the hybrid prediction head contribute complementary, non-redundant signal.

Diffusion-based generative models have recently demonstrated remarkable capacity for learning complex, multi-modal distributions across domains ranging from image synthesis to molecular generation^20–22^. LinkD-Bind leverages diffusion processes for modeling the stochastic and heterogeneous nature of molecular recognition, where binding outcomes depend on the interplay of chemical complementarity, conformational flexibility, and contextual biological factors^23^. We benchmarked LinkD-Bind against 12 methods, including three state-of-the-art deep-learning architectures (DeepDTA^7^, GraphDTA^8^, DeepPurpose^9^), a diffusion-only baseline, and conventional machine-learning approaches, on three widely used affinity datasets (BindingDB, Davis, KIBA; **Supplementary Tables S1, S2,** and **Supplementary Figs. S1, S2**). Across the nine (dataset × split) evaluation settings, LinkD-Bind ranked first in 8 of 9 by test-set RMSE and first in all three splits when averaged across datasets (**Fig. 1b**). Under random splitting, LinkD-Bind achieved RMSE values of 0.699 (BindingDB), 0.511 (Davis), and 0.447 (KIBA) with Pearson correlations of 0.862, 0.788, and 0.848, respectively, outperforming GraphDTA (RMSE = 0.800, 0.529, 0.546) and DeepPurpose (RMSE = 0.843, 0.582, 0.560) on the same benchmarks.

Performance gains were most pronounced under cold-start conditions, the settings most relevant to proteome-scale repurposing, where the majority of drug-target pairs lack experimental measurements (**Fig. 1c**). Under cold-drug splitting on BindingDB, LinkD-Bind achieved an RMSE of 1.021 versus 1.050 (GraphDTA) and 1.060 (DeepPurpose); under cold-protein splitting, 1.080 versus 1.178 and 1.203, respectively. On KIBA, where the gap was largest, LinkD-Bind reached RMSE values of 0.650 (cold drug) and 0.644 (cold protein), reducing error by 4 to 5% over the next-best model and by 6 to 17% over GraphDTA and 13 to 15% over DeepDTA across the two cold-start settings. Ablation analysis isolated the contribution of each component: removing the diffusion backbone (head only) or the hybrid prediction head (diffusion backbone only) both reduced Pearson correlation below that of full LinkD-Bind across every split, with the gaps widening under cold-protein and cold-drug evaluation (**Fig. 1c**), indicating that diffusion-based latent refinement and the hybrid head capture complementary, non-redundant signal. The sole exception was Davis cold-drug splitting, a setting with limited chemical diversity (68 unique compounds), where LinkD-Bind ranked fourth (RMSE = 0.750) but achieved the highest Pearson correlation (0.394) among similarly ranked models, indicating better rank preservation of true affinities. Together, these results establish that joint diffusion optimization over a shared drug-target latent space yields consistent improvements in both interpolation accuracy and zero-shot generalization, providing a reliable affinity foundation for downstream selectivity analysis.

### LinkD-Select enables systematic and interpretable drug selectivity analysis

A central limitation of pairwise affinity prediction is that high predicted binding to one target is uninformative about whether the same drug engages thousands of others, which may lead to unexpected adverse efects. LinkD-Select addresses this by computing selectivity from the full proteome-wide affinity distribution rather than from individual interaction scores. Across the 14,981-drug LinkD universe, predicted binding affinity is positively associated with the LinkD-Select composite score, with drugs in the high-affinity, low-entropy regime occupying the high-selectivity tail (**Fig. 2a**). Stratifying the analysis by gene functional class, comprising 298 oncogenes, 145 tumor-suppressor genes, and 36 dual-role genes, revealed that predicted binders for oncogenes attain significantly higher median selectivity than that for TSGs (Mann-Whitney U test, FDR-adjusted *P* < 0.05), consistent with their greater established druggability (**Fig. 2b**). When LinkD-Bind predictions were used to rank known drug-target interactions, a combined affinity-plus-selectivity score recovered substantially more annotated pairs than either component alone within the top 5% (**Fig. 2c**) and top 10% (**Fig. 2d**) of predictions; the combined ranking outperformed affinity-only and selectivity-only baselines across all three target classes by 5∼10 percentage points.

**Figure 2.**
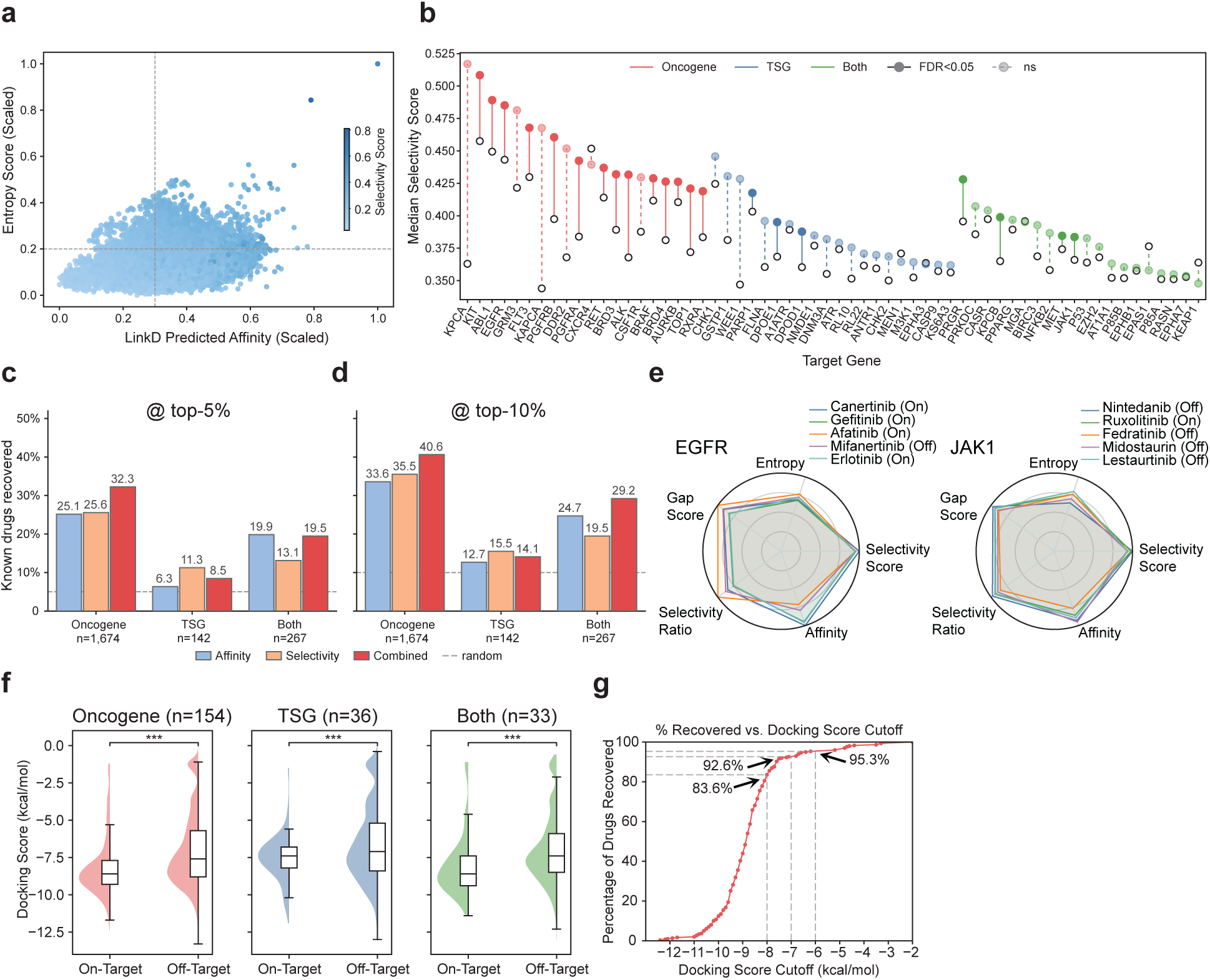
LinkD-Select enables systematic and interpretable drug-selectivity analysis across the cancer target landscape. **a.** Per-drug relationship between LinkD-Bind predicted binding affinity (x axis, scaled 0 to 1) and the Shannon entropy of the predicted-affinity distribution across 20,385 human protein targets (y axis, scaled 0 to 1) for the 14,981-drug LinkD universe; each point represents one drug, colored by its LinkD-Select composite selectivity score (legend, 0.2 to 0.8). Drugs in the upper-right quadrant exhibit high affinity, low entropy, and high selectivity, reflecting concentrated proteome-wide binding focused on a small subset of targets. **b.** Lollipop chart of median LinkD-Select score per target gene (x axis, sorted high to low) for more than 50 cancer-relevant target genes, color-coded by gene class: oncogene (red), tumor-suppressor gene (TSG; blue), or dual-role (green). Filled circles denote pairs reaching Benjamini-Hochberg FDR-adjusted *P* < 0.05 against the genome-wide background; open circles denote non-significant pairs. Oncogenes systematically attain higher median selectivity than TSGs, consistent with their established druggability. **c, d.** Fraction of known drug-target interactions recovered within the top 5% (**c**) and top 10% (**d**) of LinkD predictions, stratified by target class (oncogene, *n* = 1,674; TSG, *n* = 142; both, *n* = 267). Three ranking strategies are compared: affinity-only (blue), selectivity-only (orange), and a combined score (red), with the random-recovery baseline shown as a dashed line. The combined affinity-plus-selectivity ranking yields the best recovery in each class, exceeding either component alone by 5 to 10 percentage points. **e.** Radar plots for two representative oncogenic targets, EGFR (left) and JAK1 (right), comparing five compounds each across five dimensions: Selectivity Score, Entropy, Gap Score, Selectivity Ratio, and Affinity. Compounds known to engage the target on-target (for example, Erlotinib for EGFR; Ruxolitinib for JAK1) display coordinated profiles of high affinity, large affinity gap, and elevated composite score, whereas off-target compounds yield flattened profiles. **f.** Molecular-docking score distributions for on-target versus off-target drug-target pairs, stratified by target class (oncogene, *n* = 154; TSG, *n* = 36; both, *n* = 33). On-target interactions are significantly more favorable than off-target interactions in every class (Mann-Whitney *U* test, *** *P* < 0.001). **g.** Cumulative recovery of known drug-target interactions as the docking-score cutoff is relaxed from −12 to −2 kcal/mol. At cutoffs of −8.0, −7.0, and −6.0 kcal/mol, 83.6%, 92.6%, and 95.3% of annotated pairs are recovered respectively, demonstrating that LinkD-Select prioritized predictions are enriched among structurally plausible interactions.

At the individual target level, LindD-Select radar profiles for EGFR and JAK1 show that selective on-target compounds (Erlotinib, Ruxolitinib) display coordinated high values across affinity, gap score, and selectivity ratio, whereas off-target compounds yield flattened profiles with minimal affinity separation (**Fig. 2e**). To assess whether these computationally derived selectivity patterns correspond to physically meaningful binding differences, we performed molecular docking for annotated drug-target pairs. On-target docking scores were significantly more favorable than off-target scores across oncogenes, TSGs, and dual-role targets (Mann-Whitney *U* test, *P* < 0.001; **Fig. 2f**), providing orthogonal validation using structure-based approaches. Cumulative recovery analysis demonstrated that 95.3% of known drug-target pairs were recovered at a docking-score cutoff of −6.0 kcal/mol, with rapid saturation as the threshold became more permissive (**Fig. 2g**), confirming that LinkD-Select prioritizes interactions independently supported by structural evidence. Together, these results establish that proteome-wide selectivity, quantified as a distributional property of predicted affinity rather than a single pairwise estimate, provides a principled and scalable metric for distinguishing on-target from off-target interactions across diverse target classes.

### LinkD-Pheno integrates cellular perturbation data to validate drug selectivity

To test whether LinkD-predicted selectivity translates into functional cellular phenotypes, we integrated large-scale drug-perturbation and CRISPR-Cas9 gene-knockout datasets across 960 cancer cell lines spanning 13 cancer tissue lineages (**Fig. 3a**). Drug-response AUC profiles for 6,532 compounds and Chronos gene-dependency scores (DepMap 25Q1) for 7,502 genes were jointly analyzed, with functional concordance defined as the Pearson correlation between drug sensitivity and gene dependency across shared cell lines (median 327 per pair, range 15 to 605). Multiple-testing correction used the Benjamini-Hochberg procedure with adjusted *P* < 0.05, yielding 464,820 evaluable drug-gene records. Tissue representation was heaviest in lung (178 cell lines), blood (170), and urogenital cancers (104), and lightest in soft tissue (21) and thyroid (16) lineages (**Fig. 3b**), reflecting screening coverage and tissue-specific target availability.

**Figure 3.**
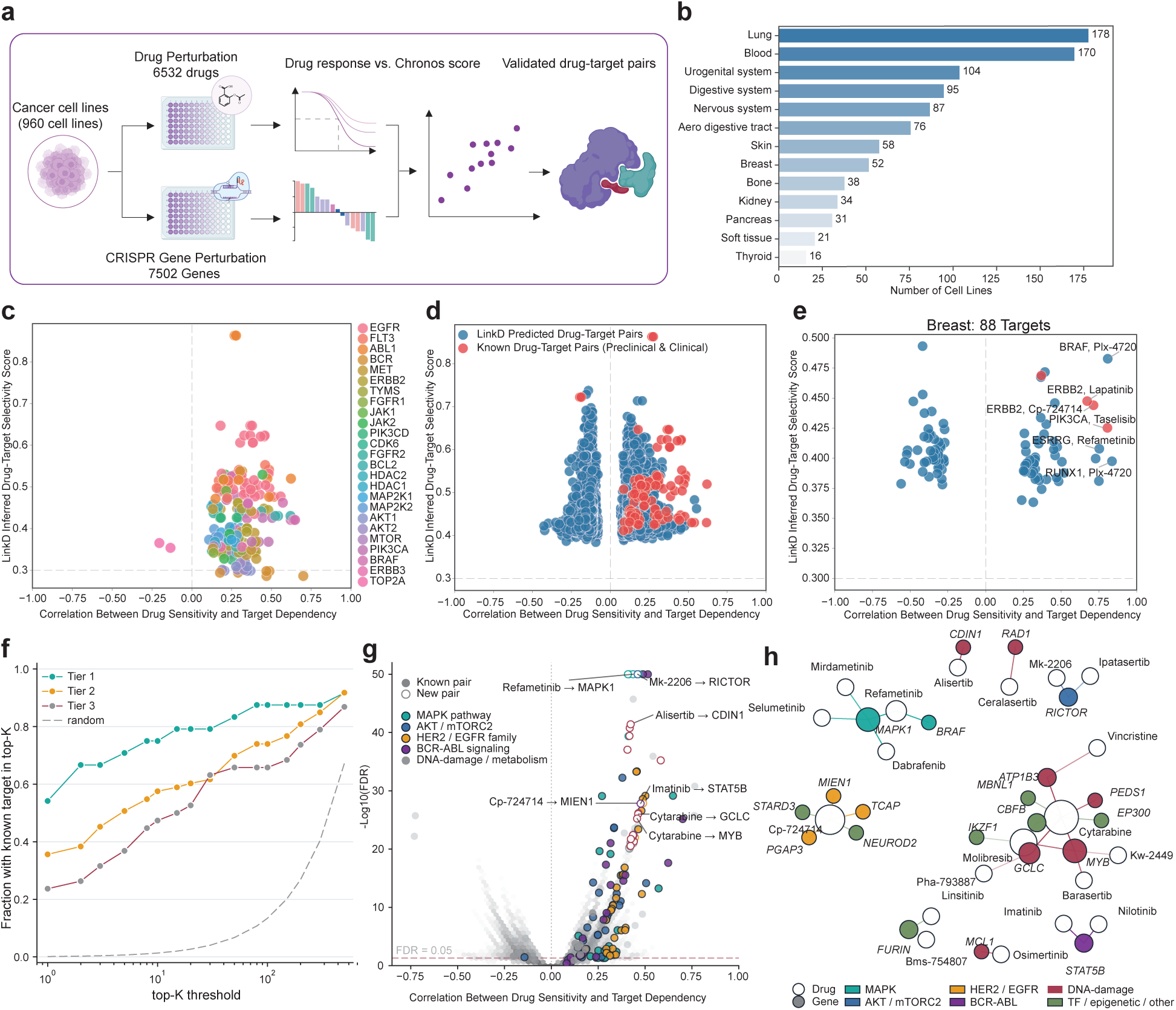
Phenotypic validation of LinkD-Select drug-selectivity predictions using large-scale drug-perturbation and CRISPR-dependency data. **a.** Schematic of the LinkD-Pheno validation framework. Drug-response AUC profiles for 6,532 compounds and CRISPR-Cas9 gene-dependency Chronos scores for 7,502 genes were intersected across 960 cancer cell lines. For each drug-gene pair, functional concordance was defined as the Pearson correlation between drug sensitivity and gene dependency across shared cell lines; statistically significant concordant pairs define the validated drug-target set. **b.** Distribution of cell lines across 13 cancer tissue lineages. **c.** Scatter of phenotypic concordance (x axis: Pearson correlation between drug sensitivity and gene dependency) versus LinkD-Select score (y axis) for drug-target pairs whose target is one of 25 canonical oncology genes. Selective pairs concentrate in the upper-right quadrant, indicating positive concordance and high selectivity. **d.** Same axes extended to all LinkD-predicted drug-target pairs (blue), with ChEMBL-annotated preclinical and clinical reference pairs overlaid (red). Known pairs occupy the same upper-right regime as the broader predicted set, demonstrating that LinkD novel predictions do not lie in random-noise tails. **e.** Tissue-specific recovery in breast cancer: scatter restricted to the 88 LinkD-prioritized targets evaluated across 52 breast cancer cell lines. Labeled points include canonical known drug-target and novel candidates within the same selectivity-concordance regime. **f.** Cumulative fraction of LinkD-predicted drug-target pairs whose ChEMBL-annotated target is recovered within the top *K* concordance-ranked candidates per drug, stratified into three LinkD-Select confidence tiers. The grey dashed line indicates the uniform-random baseline. Tier 1 achieves approximately 85% recovery by *K* = 50. **g.** Pathway-classified discovery volcano plot. **h.** Discovery network of the 34 novel pairs.

Across 25 canonical oncology targets, LinkD-prioritized pairs concentrated in the high-selectivity, high-concordance regime (**Fig. 3c**), and ChEMBL-annotated reference pairs occupied the same upper-right region as the broader LinkD-predicted set (**Fig. 3d**), demonstrating that LinkD predictions are not random-noise candidates but co-localize with clinically validated drug-target relationships. Tissue-specific analysis in breast cancer (88 LinkD-prioritized targets, 52 cell lines) recovered known drug-target pairs (BRAF and Plx-4720; ERBB2 and Lapatinib; ERBB2 and Cp-724714; PIK3CA and Taselisib) and surfaced novel breast-cancer candidates (RUNX1 and Plx-4720; ESRRG and Refametinib) within the same regime (**Fig. 3e**). When drugs were stratified into LinkD-Select quartiles, top-*K* recovery followed a monotonic ordering, with the top quartile reaching approximately 85% annotated-target recovery within the top 50 concordance-ranked candidates per drug, an order of magnitude above the uniform-random baseline (**Fig. 3f**).

A pathway-classified discovery analysis identified 211 ChEMBL-annotated drug-target pairs and 34 novel pairs passing the stringent thresholds |ρ| ≥ 0.40 and −log_10_(FDR) ≥ 20 (**Fig. 3g**); annotated and novel discoveries co-clustered by pathway (MAPK, AKT/mTORC2, HER2/EGFR family, BCR-ABL signaling, DNA damage/metabolism), demonstrating that LinkD novel predictions extend along the same biological axes as validated reference pairs rather than scattering into noise. Projecting the 34 novel pairs onto a drug-gene network revealed three biologically coherent structures (**Fig. 3h**): cytarabine emerged as the largest discovery hub with seven novel partners spanning transcription-factor, metabolic, and epigenetic genes (MYB, GCLC, EP300, CBFB, MBNL1, ATP1B3, PEDS1); Cp-724714 formed a tight 17q12 amplicon module engaging five co-amplified genes (MIEN1, PGAP3, TCAP, STARD3, NEUROD2); and three MEK inhibitors (Refametinib, Mirdametinib, Selumetinib) converged on the downstream node MAPK1, providing within-class consistency. Generalization across the eight cancer tissue lineages further confirmed these patterns (**Supplementary Figs. S3, S4**), with selective drug-target pairs consistently localized in regions of high predicted affinity and positive drug-dependency concordance regardless of tissue context.

### Real-world evidence validation of LinkD predictions using population-scale EHR

To assess whether LinkD-predicted drug, target, and disease relationships extend beyond molecular and cellular phenotypes to real-world clinical outcomes, we performed population-scale validation using longitudinal electronic health record (HER) data from two independent cohorts totaling 11.5 million individuals (**Fig. 4a**, see Methods). LinkD-supported drug and disease associations formed coherent networks connecting drugs, targets, and cancer phenotypes (**Fig. 4b**), with candidate associations contextualized by LinkD-predicted drug and target interactions, as well as gene and disease association scores from Open Targets across 50 cancer types. Several predicted associations converged on well-characterized oncogenic pathways, while others suggested repositioning opportunities supported by both molecular selectivity scores and EHR-derived risk estimates. Cancer-focused scans in both the Mount Sinai Data Warehouse (MSDW) EHR data identified 718 unique drug-cancer pairs across 286 drugs and 55 ICD-10 cancer codes, with 91.1% reaching nominal significance at *P* < 0.05 and protective effects against disease occurrence (median OR = 0.49, range 0.061–0.895; **Supplementary Table S3**). Analyses in the UKB cohort found 241 unique drug-cancer pairs across 143 drugs and 20 ICD-10 cancer codes (median OR = 0.21, range 0.003–1.301; **Supplementary Table S4**). These results showed that drugs prioritized by LinkD to bind strong cancer targets are enriched among associations with reduced cancer risk, as reflected by odds ratios below unity and increased statistical significance relative to background comparisons (**Supplementary Fig. S5a,b**). The most frequently represented cancer types were consistent across both cohorts, with prostate cancer (C61), thyroid cancer (C73), non-melanoma skin cancer (C44), breast cancer (C50), and liver cancer (C22) accounting for the largest numbers of target-drug associations in both datasets. All 241 UKB cancer drug-disease pairs were also identified in the MSDW analysis, providing independent replication across two demographically distinct populations.

**Figure 4.**
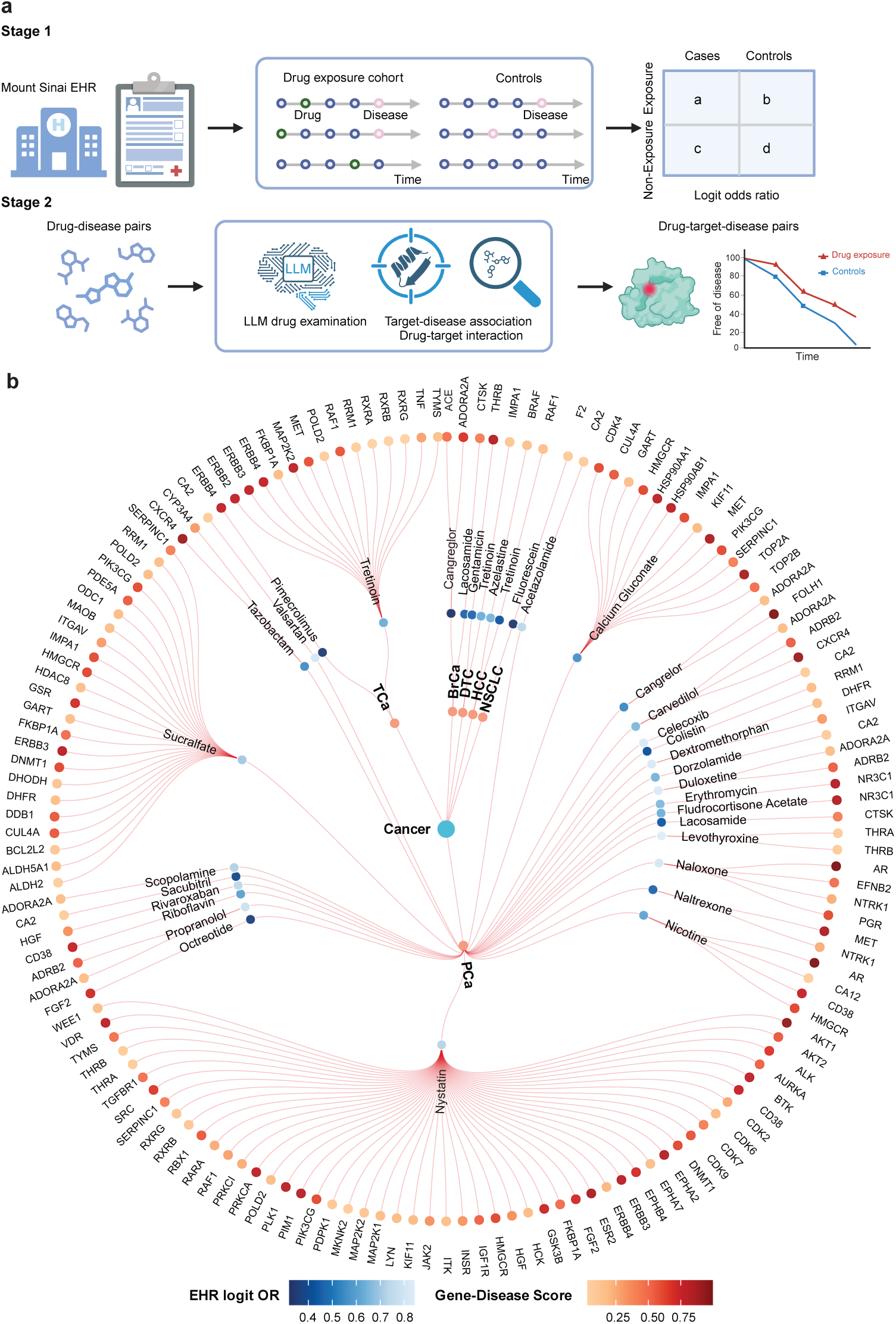
Population-scale real-world evidence validation of LinkD predictions. **a.** Overview of the EHR-based validation framework integrating drug exposure, disease outcomes, LinkD-predicted drug and target interactions, and gene-disease associations across two independent cohorts (Mount Sinai Health System and UK Biobank, totaling 11.5 million individuals). **b.** Network representation of drug-target-disease associations supported by both LinkD predictions and EHR-derived risk estimates. Drug-disease edges and disease nodes are colored by EHR logit odds ratio; gene-disease edges and gene nodes are colored by Open Targets gene-disease association score. Odds ratios are estimated using propensity-score-matched logistic regression.

Detailed analyses of representative drug-disease pairs demonstrated reproducible protective associations, azelastine and liver cancer (**Supplementary Fig. S5c**; summary OR = 0.69, *P* < 0.001) and tretinoin and thyroid cancer (**Supplementary Fig. S5d**; summary OR = 0.43, *P* < 0.001), each of which yielded odds ratios consistently below unity across ten matched sensitivity analyses. These examples illustrate how the LinkD framework can integrate evidence from molecular interaction modeling through phenotypic prioritization and be validated with population-level observational evidence, with each layer providing independent support for the same drug-disease hypotheses. Although these EHR analyses are observational and do not establish causality, the consistency of protective directionality across independent cohorts, sensitivity specifications, and permutation controls suggests that LinkD-prioritized compounds are enriched for clinically relevant signals that warrant prospective evaluation.

### Experimental validation of β-blocker-ADRB2 interactions in prostate cancer

To evaluate LinkD end-to-end in a disease-relevant setting, we focused on prostate cancer and selected ADRB2 binders as a high-priority repurposing candidates based on convergent evidence across LinkD modules: high predicted affinity and selectivity (LinkD-Bind and LinkD-Select), concordant drug-sensitivity and gene-dependency signals (LinkD-Pheno), and protective EHR associations (LinkD population-scale validation). Among the 14,981-drug universe, propranolol (rank 1) and carvedilol (rank 3) emerged as the top-selectivity candidates for ADRB2, while the β1-selective metoprolol scored substantially lower (**Fig. 5a**), providing a within-class negative-control comparator. We conducted complementary structural, cell-based, and real-world validations centered on the androgen-sensitive prostate cancer cell line LNCaP.

**Figure 5.**
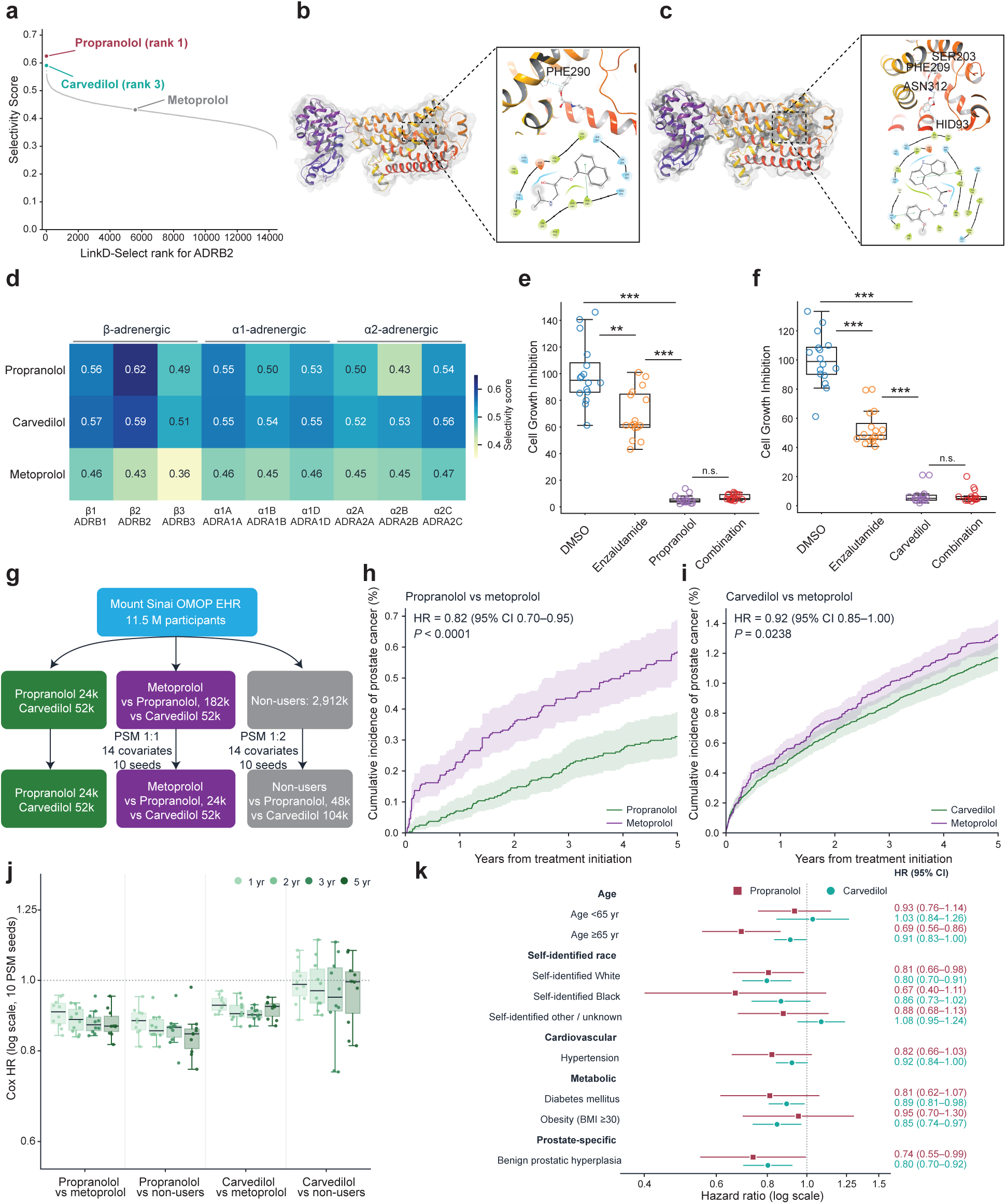
Mechanistic, cellular, and population-scale validation of LinkD-prioritized β-blocker, ADRB2, and prostate cancer associations. **a.** LinkD-Select rank of all 14,981 drugs against ADRB2, with rank position (x axis) plotted against per-drug selectivity score (y axis). Propranolol (rank 1; red), carvedilol (rank 3; teal), and metoprolol (grey) are labeled. Propranolol and carvedilol emerge as the highest-priority candidates, while metoprolol, the β1-selective β-blocker, scores substantially lower. **b, c.** Receptor-centric molecular docking of propranolol (**b**; AutoDock Vina score −8.0 kcal/mol) and carvedilol (**c**; −9.9 kcal/mol) into the canonical transmembrane binding pocket of ADRB2 (PDB 2RH1). Cartoon depictions show the full receptor with the bound ligand; insets show the ligand-binding pocket with annotated stabilizing contacts (representative residues PHE-290 and HIS-93 are labeled along with conserved transmembrane residues). **d.** Drug-by-receptor selectivity heatmap (LinkD-Select score per cell) for propranolol, carvedilol, and metoprolol against the nine adrenergic-receptor subtypes (β-adrenergic, ADRB1 to ADRB3; α1-adrenergic, ADRA1A, ADRA1B, ADRA1D; α2-adrenergic, ADRA2A to ADRA2C). Propranolol and carvedilol attain the highest scores at ADRB2, whereas metoprolol scores substantially lower across the panel. **e, f.** Growth-inhibition assays in the androgen-sensitive prostate cancer cell line LNCaP after 72 h treatment with DMSO control, enzalutamide (AR antagonist, 10 µM), propranolol (100 µM; **e**) or carvedilol (50 µM; **f**), and the corresponding two-drug combinations. Boxes show the interquartile range; whiskers extend to the 5th and 95th percentiles; points represent individual wells (biological duplicates, 8 wells per condition). Both β-blockers reduce LNCaP viability more strongly than enzalutamide alone; combining either β-blocker with enzalutamide does not further reduce growth (N.S.). Statistical significance is denoted as \*\**P* < 0.01, \*\*\**P* < 0.001 (Mann-Whitney *U* test with Bonferroni correction). **g.** Mount Sinai OMOP-mapped EHR cohort (11.5 million participants) propensity-score-matching design. Propranolol-exposed, carvedilol-exposed, and metoprolol-exposed patients are matched 1:1 within the β-blocker class and 1:2 against non-users on 14 demographic, comorbidity, and prescription covariates. **h, i.** Cumulative incidence of prostate cancer over 5 years from treatment initiation, comparing propranolol versus metoprolol (**h**; HR = 0.82, 95% CI 0.70 to 0.95, *P* < 0.0001) and carvedilol versus metoprolol (**i**; HR = 0.92, 95% CI 0.85 to 1.00, *P* = 0.0238). Both β-blockers are associated with lower prostate cancer incidence relative to metoprolol, the within-class negative-control comparator. **j.** Cox hazard ratios across four pairwise comparisons (propranolol versus metoprolol; propranolol versus non-users; carvedilol versus metoprolol; carvedilol versus non-users) at four follow-up windows (1, 2, 3, and 5 years). Hazard ratios remain consistently below 1, demonstrating temporal stability of the protective effect. **k.** Subgroup hazard-ratio forest plot for propranolol (red) and carvedilol (teal) stratified by age, self-identified race, cardiovascular comorbidity (hypertension), metabolic comorbidity (diabetes mellitus, obesity, BMI ≥ 30), and prostate-specific covariate (benign prostatic hyperplasia). Protective effects are consistent across all subgroups examined.

At the structural level, receptor-centric docking of propranolol (−8.0 kcal/mol) and carvedilol (−9.9 kcal/mol) into ADRB2 (PDB 2RH1) yielded predicted poses localized to the canonical transmembrane pocket (**Fig. 5b, c**), with stabilizing contacts consistent with established β-adrenergic recognition motifs. Cross-receptor LinkD-Select profiling across the nine adrenergic-receptor subtypes confirmed that propranolol and carvedilol attain their highest scores at ADRB2, whereas metoprolol scores uniformly lower (**Fig. 5d**). To assess functional consequences, LNCaP cells were treated for 72 h with propranolol (100 µM) or carvedilol (50 µM), enzalutamide (AR antagonist, 10 µM), or the corresponding two-drug combinations (biological duplicates, 8 wells per condition). Both β-blockers elicited stronger growth inhibition than enzalutamide alone (**Fig. 5e, f**); combining either β-blocker with enzalutamide did not further reduce growth (N.S.), suggesting that β2-adrenergic blockade dominates the antiproliferative phenotype under these conditions, or that adrenergic and AR signaling converge on a shared limiting process.

Population-level evidence from the MSDW EHR cohort corroborated these findings. Within an 11.5-million-participant cohort, propranolol-exposed, carvedilol-exposed, and metoprolol-exposed patients were propensity-score matched 1:1 within class and 1:2 against non-users on 14 demographic and clinical covariates (**Fig. 5g**). Over a 5-year follow-up, propranolol was associated with a reduced incidence of prostate cancer relative to metoprolol (HR = 0.82, 95% CI 0.70 to 0.95, *P* < 0.0001; **Fig. 5h**), as was carvedilol (HR = 0.92, 95% CI 0.85 to 1.00, *P* = 0.0238; **Fig. 5i**). Protective hazard ratios remained stable across follow-up windows of 1, 2, 3, and 5 years (**Fig. 5j**), and were consistent across age strata, self-identified race, cardiovascular and metabolic comorbidities, and benign prostatic hyperplasia (**Fig. 5k**). Although these EHR associations are observational and do not establish causality, the consistency of protective directionality across independent comparisons, follow-up windows, and clinical subgroups, in combination with the structural docking and LNCaP growth-inhibition evidence, provides multi-layer support for the ADRB2 repurposing hypothesis and illustrates the LinkD framework’s capacity to triangulate predictions across mechanistic, cellular, and population scales.

### LinkD-Agent integrates mechanistic, phenotypic, and clinical evidence for interactive drug repurposing

To translate LinkD predictions into an end-to-end repurposing platform, we developed LinkD-Agent, an interactive web-based AI system (https://linkd-agent.onrender.com/) that executes natural-language queries as auditable, multi-step analyses that integrate mechanistic, functional, and clinical evidence (**Fig. 6**). The agent is built on a structured knowledge base comprising 276,147 drug, target, and disease associations from clinical trial records (4,274 drugs, 1,520 targets, 2,684 diseases), 13,008 curated causal gene and disease relationships (3,400 genes, 3,859 diseases), and functional annotations for 1,029 oncogenes and tumor suppressor genes (485 oncogenes, 404 tumor suppressors, 140 dual-role). This deterministic data layer is integrated with all upstream LinkD outputs: binding affinity and selectivity profiles for 14,981 drugs across 20,385 human protein targets, 464,820 drug and gene dependency correlation records, and EHR-derived association statistics from the MSDW (41,120 associations across 730 drugs) and UK Biobank (693 associations across 143 drugs) real-world cohorts.

**Figure 6.**
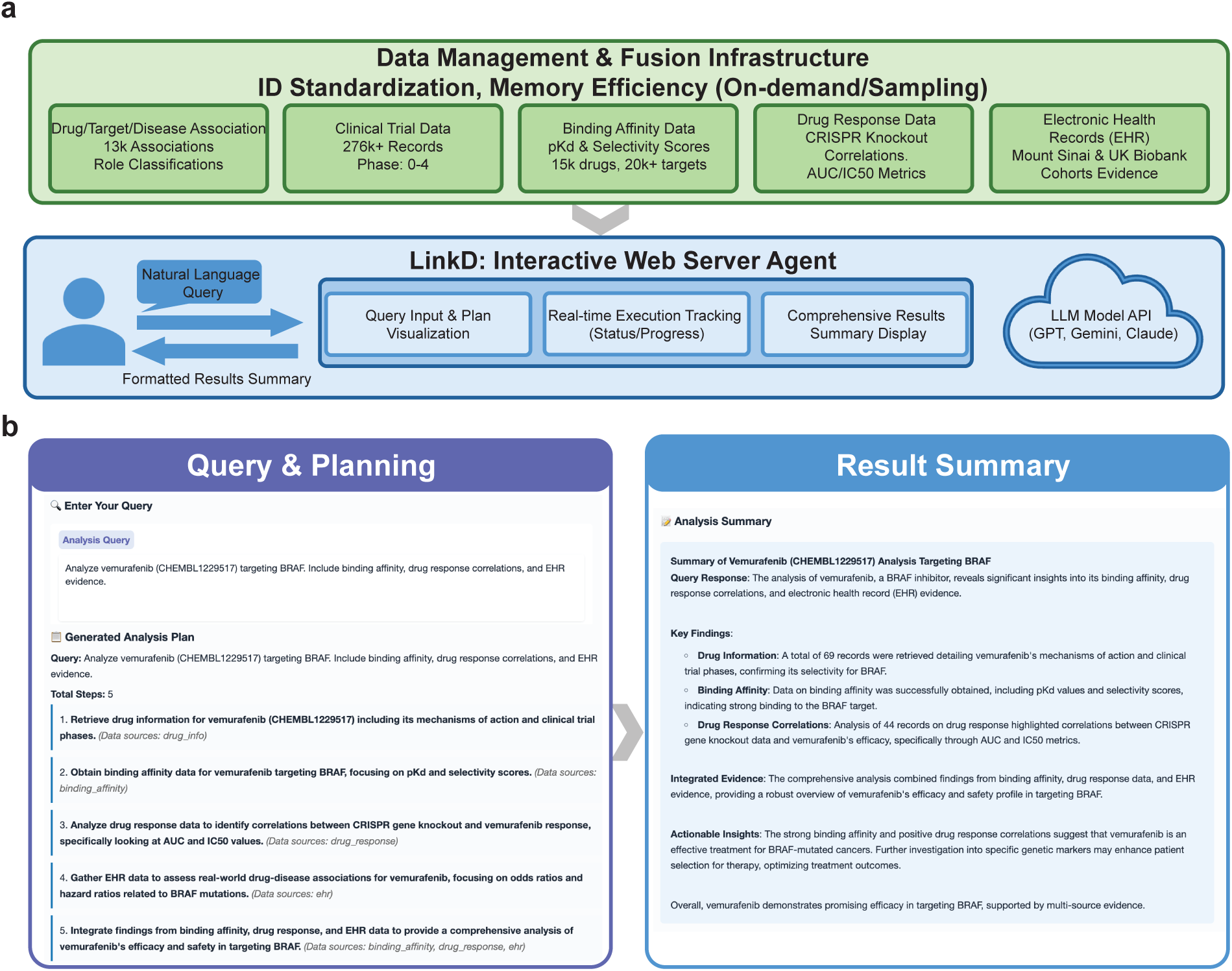
Interactive AI agent for integrated drug repurposing analysis. **a.** System architecture of the LinkD AI agent, exposing modular tools for drug selectivity, gene dependency, and EHR-derived drug effect querying through a Model Context Protocol server. **b.** Example of agent planning, in which a natural-language query is decomposed into structured function calls and execution steps. Integrated output summarizing mechanistic evidence, oncogene dependency signals, EHR protective effect strength, and an overall drug repurposing score.

The agent exposes these resources through a Model Context Protocol (MCP) tool interface coupled to a large language model reasoning layer (supporting GPT, Gemini, and Claude LLMs) that separates deterministic data retrieval from natural-language interpretation (**Fig. 6a**). Critically, all numerical statistics are derived directly from structured database queries, while the language model is used solely for query parsing, plan generation, and result synthesis. Upon receiving a query, the agent extracts biomedical entities (drug identifiers, gene symbols, disease names, ICD codes), decomposes complex requests into sequential analytical steps, and orchestrates a chain of backend calls: typically retrieving selective targets from LinkD affinity and selectivity profiles, prioritizing targets with concordant functional dependencies, and contextualizing candidates with clinical trial and EHR evidence (**Fig. 6b**). Real-time execution tracking exposes each reasoning step, making the analytical process fully inspectable and reproducible.

Outputs are returned as structured summaries that consolidate multi-source evidence into reports suitable for hypothesis generation. For example, querying a candidate drug returns its top selective targets with pKd values and selectivity categories, concordant CRISPR dependency signals with effect sizes, relevant clinical trial phases, and EHR-derived odds ratios with confidence intervals, all within a single integrated report. This design ensures that LinkD-Agent functions not as a black-box recommender but as a transparent evidence aggregation system, enabling domain experts to evaluate the strength and consistency of repurposing hypotheses across independent data modalities before committing to experimental or clinical follow-up.

Beyond standard query-triggered evidence aggregation, LinkD-Agent extends to an autonomous virtual-clinical-trial mode that operationalizes target-trial emulation directly from OMOP-mapped EHR data (**Supplementary Fig. S6**). A natural-language clinical question is decomposed into six sequential modules: protocol designer, cohort builder, three-arm trial constructor (drug versus within-class active comparator versus untreated control), causal estimator with propensity-score matching and average treatment effect on the treated, robustness and reporting (sensitivity analyses, negative controls, prespecified subgroups), and final evidence report. This pipeline produces auditable hazard-ratio or odds-ratio estimates with forest-plot summaries, enabling users to test a LinkD repurposing hypothesis using a transparent, prespecified observational analysis based on real-world data.

## Discussion

LinkD bridges the longstanding divide between target-centric and phenotype-driven drug repurposing by jointly modeling molecular interactions, cellular perturbation phenotypes, and real-world clinical evidence within a unified diffusion-based architecture. This multi-scale integration delivers substantial gains in drug-target affinity prediction, enables systematic and interpretable selectivity analysis, and produces repurposing hypotheses corroborated by molecular, cellular, and population-level evidence.

A key conceptual advance of LinkD is the shift from isolated pairwise affinity prediction to distributional selectivity analysis. Most existing deep learning approaches for drug-target interaction modeling evaluate each drug-target pair independently, producing a single affinity score devoid of proteome-wide context^7–9,24^. While such scores are useful for ranking candidate binders, they provide limited insight into whether a compound engages its intended target selectively or interacts promiscuously across the proteome—a distinction with profound implications for therapeutic efficacy, toxicity, and clinical translatability^10,25^. By computing entropy over the predicted affinity distribution and defining selectivity as a function of relative binding preference, LinkD enables direct quantification of target specificity. Our analyses revealed that this entropy-aware framework recapitulates known differences in druggability between oncogenes and tumor suppressor genes and identifies selective interactions that are validated by both molecular docking and phenotypic perturbation data, establishing distributional selectivity as a tractable and biologically meaningful metric for computational drug repurposing.

The integration of large-scale phenotypic evidence constitutes a second distinguishing feature of LinkD. By jointly analyzing pharmacological sensitivity profiles across hundreds of cancer cell lines with CRISPR-based genetic dependency maps, we established a phenotype-based validation layer that complements and extends structure-informed predictions. The consistent enrichment of known drug-target pairs among high-selectivity, high-concordance predictions across diverse tissue lineages supports the hypothesis that selective molecular binding translates into functional cellular vulnerability—a relationship that is assumed but rarely demonstrated at this scale. Importantly, this phenotypic validation is not limited to canonical oncology targets; the tissue-specific generalization analyses (**Supplementary Figs. S3 and S4**) suggest that the framework captures biologically coherent drug-target-tissue relationships beyond well-studied receptor tyrosine kinases, indicating broad applicability across cancer biology.

The translation of computational predictions to real-world clinical evidence represents perhaps the most consequential contribution of this work. The majority of published repurposing frameworks conclude their validation at the molecular or cellular level, leaving a critical translational gap between predicted interactions and observable clinical outcomes^17,26^. By leveraging longitudinal EHR data from two independent health systems—the Mount Sinai Health System and UK Biobank—we demonstrated that LinkD-prioritized drug-disease associations are enriched among protective clinical signals, with effect directions that are broadly consistent across cohorts. The experimental validation of β-blocker-ADRB2 interactions in prostate cancer further illustrates how LinkD predictions can be triangulated across evidence scales: from structural modeling of receptor engagement, through functional growth suppression in LNCaP cells, to population-level associations with reduced prostate cancer risk. While we emphasize that these EHR findings are observational and do not establish causality, they demonstrate that structure- and phenotype-informed selectivity signals can be reflected at the population level, substantially strengthening the evidentiary basis for prioritized repurposing hypotheses.

From a methodological perspective, the application of diffusion-based generative models to drug-target interaction prediction offers several advantages over conventional discriminative architectures. The diffusion framework naturally accommodates the stochastic and multi-modal nature of molecular recognition, where binding outcomes depend on a complex interplay of chemical complementarity, conformational dynamics, and contextual biological factors^20–22^. The superior generalization of LinkD under cold-drug and cold-protein evaluation protocols suggests that the diffusion process learns transferable representations of binding determinants that extend beyond memorized chemical or sequence patterns. This property is particularly valuable for drug repurposing, where the most clinically impactful discoveries often involve novel drug-target combinations that lie outside the training distribution of existing models.

The deployment of LinkD as an interactive AI agent addresses a practical barrier that limits the adoption of complex computational repurposing frameworks by domain experts. By encapsulating multi-modal evidence retrieval, integration, and interpretation within a natural-language interface, the agent enables clinicians and biologists to interrogate repurposing hypotheses without specialized computational skills. The modular architecture, which couples a data management layer with a language-model-driven tool orchestration system, ensures that analyses are transparent, auditable, and reproducible—qualities that are essential for generating hypotheses that will ultimately require prospective clinical evaluation.

Several limitations of this study warrant discussion. First, although LinkD integrates multiple evidence modalities, the framework relies on the quality and coverage of available training data. Drug-target interaction databases are biased toward well-studied targets and compound classes^27^, and phenotypic screening panels, while expansive, do not comprehensively cover the druggable genome. These data biases may limit the framework’s ability to identify interactions involving understudied targets or chemically novel compounds. Second, the EHR-based validation, while providing valuable population-level context, is inherently observational and subject to confounding, indication bias, and incomplete ascertainment of drug exposures and clinical outcomes^18,19^. The associations reported here should therefore be interpreted as hypothesis-generating rather than confirmatory, and will require prospective validation in appropriately designed clinical studies. Third, the experimental validation was conducted in a single cell line (LNCaP), and the generalizability of the observed anti-proliferative effects to other prostate cancer subtypes and *in vivo* models remains to be established. Fourth, the current implementation of the AI agent depends on external LLM APIs, which introduces considerations around reproducibility, cost, and data governance that will need to be addressed for clinical deployment.

Looking forward, several extensions of LinkD could further enhance its utility. Incorporating three-dimensional protein structure information—now widely available through AlphaFold^28^ and related methods—could improve the resolution of binding-site modeling and enable structure-guided selectivity optimization. Integration of additional phenotypic modalities, such as single-cell transcriptomic responses to drug perturbation^29^ or spatial transcriptomics of treated tissues, could further refine the phenotype-based validation layer. The EHR validation framework could be extended to incorporate time-to-event analyses, propensity score matching, and target trial emulation to strengthen causal inference from observational data^30^. Finally, the interactive agent could be enhanced with active learning capabilities, enabling it to propose optimally informative experiments for validating high-priority repurposing candidates.

In summary, LinkDLinkD establishes a new paradigm for computational drug repurposing that integrates target-centric molecular modeling with phenotypic validation and real-world clinical evidence within a unified framework. By operating across molecular, cellular, and population scales, LinkD addresses a critical gap in existing repurposing approaches and provides a principled approach for generating, prioritizing, and stress-testing drug repurposing hypotheses with converging evidence at every scale.

## Materials and Methods

### Drug-target interaction and binding affinity data

The annotated drug-target interaction data were collected from publicly available pharmacological resources, including BindingDB (2025-03)^31^, and from benchmark binding affinity datasets (Davis and KIBA from Therapeutics Data Commons^32^, https://tdcommons.ai/) that report experimentally measured dissociation constants, inhibition constants, or activity values. Drug molecular structures were obtained as SMILES strings and standardized using canonicalization to remove salts and normalize atom representations. Protein sequences were retrieved from reference proteomes, with isoforms collapsed to a representative canonical sequence when necessary. Only drug-target pairs with unambiguous target mapping and valid chemical structures were retained. Binding affinity values were log-transformed where appropriate to harmonize measurement scales across sources. For BindingDB and Davis dataset, we used the dissociation constant *K*_*d*_ in which affinities are reported in nanomolar (nM) units. The raw *K*_*d*_ values were converted to *pK*_*d*_ = −log _10_(*K*_*d*_ ∗ 10^−9^). On this scale, higher values denote stronger binding affinity. For the KIBA dataset, binding affinities are expressed as KIBA scores, a dimensionless measure that combines Kd, Ki, and IC_50_ measurements by optimizing their statistical consistency. No logarithmic transformation has been applied to the KIBA scores. Descriptive statistics for the datasets are provided in Supplementary Table S1.

### Molecular Feature Embedding and Extraction

Small-molecule drugs are represented by SMILES strings and encoded using ChemBERTa^33^, a pretrained Transformer model for chemical language modeling. Given a drug SMILES string *s*_*d*_for drug *d*, the model produces a contextualized embedding with a vector size *D*_*d*_ of 768,

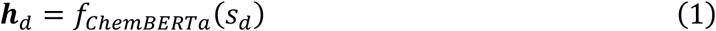

where 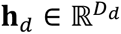 denotes the learned drug embedding.

Protein targets are represented by their amino-acid sequences and encoded using the pretrained protein language model ESM2^34^. For a protein sequence *s*_*t*_, ESM2 generates residue-level embeddings, which are aggregated to obtain a 1024 fixed-length target representation:

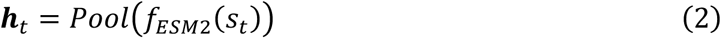

where Pool denotes mean aggregation across residues unless otherwise specified. Drug and target embeddings are projected into a shared interaction space using learned linear transformations

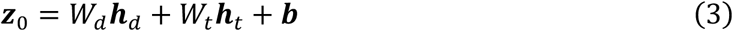

where 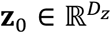 represents the initial joint drug-target latent embedding.

### Diffusion-based training process

LinkD models drug-target interactions using a denoising diffusion probabilistic model (DDPM). In the forward diffusion process, Gaussian noise is added to the joint latent embedding,

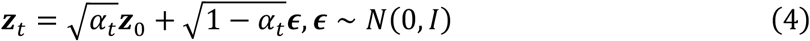

where *t* ∈ {1, …, *T*} denotes the diffusion step and *α*_*t*_ follows a predefined noise schedule. A neural network *ϵ*_*θ*_ is trained to predict the added noise,

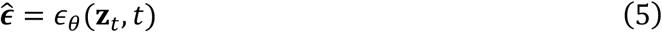

allowing recovery of the clean latent representation,

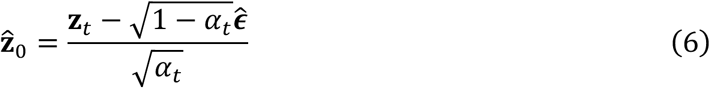

The diffusion loss is defined as,

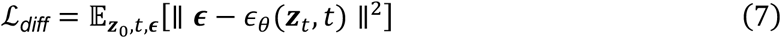

The recovered latent embedding **ẑ**_0_ is passed to a prediction head consisting of a multilayer perceptron followed by a Random Forest regressor.

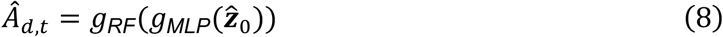

where *Â_*d*,*t*_* denotes the predicted binding affinity.

The total training objective combines diffusion loss and supervised affinity loss:

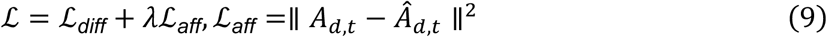

Models are optimized using the Adam optimizer with a fixed learning rate and early stopping based on the validation loss.

### Model training and evaluation

LinkD was benchmarked against established drug-target affinity prediction methods following standard evaluation protocols used for the BindingDB, Davis, and KIBA datasets. Comparator models included DeepPurpose^9^, GraphDTA^8^, DeepDTA^7^, baseline diffusion model, and classical baselines (linear regression, support vector regression, random forest, gradient boosting, XGBoost, multilayer perceptron, feedforward neural networks, graph convolutional networks, and deep cross networks). All models were trained and evaluated using identical dataset splits, input representations, and target definitions to ensure fair comparison. Each dataset was divided into training, validation, and test subsets in an 8:1:1 ratio. Three splitting methodologies were employed: random, in which drug-target pairs were assigned at random; cold-drug, in which all test-set compounds were excluded from training and validation; and cold-protein, in which all test-set proteins were excluded from training and validation. The cold-drug and cold-protein paradigms assess zero-shot generalization, necessitating the model to predict affinities for entities entirely absent from the training data.

Model performance was assessed on held-out test sets using standard regression metrics: root mean squared error (RMSE), mean squared error (MSE), mean absolute error (MAE), median absolute error (MdAE), coefficient of determination (R^2^), and Pearson correlation coefficient (r) between predicted and measured affinities. Lower RMSE, MSE, MAE, and MdAE indicate improved accuracy, whereas higher R^2^ and Pearson r indicate stronger explanatory power and rank consistency. For each dataset and split setting, models were ranked using a multi-metric scheme: for each metric, models were ranked independently (lower is better for error metrics; higher for correlation-based metrics), and ranks were aggregated to yield an overall performance ranking. Rankings were further summarized across split settings to assess robustness under distribution shift. Results correspond to mean performance over five independent runs with different random seeds.

### Drug- and target-centric selectivity scoring

LinkD quantifies selectivity using complementary drug-centric and target-centric formulations to characterize the specificity of predicted drug-target interactions at the proteome scale. Binding affinities are predicted by LinkD and expressed in *pK*_*d*_-like units, denoted *Â_d,t_*, for drug *d*and protein target *t*. The pan-proteome target panel comprises 20,385 human protein targets assembled from UniProt reference proteomes; binding affinity was predicted for every drug-target combination in the universe (14,981 drugs × 20,385 targets = 305,387,685 pairs), and this panel is held constant during evaluation to ensure consistent normalization of distribution-based selectivity measures. Drug-centric selectivity describes how concentrated a drug’s predicted binding profile is across the proteome. For each drug *d*, predicted affinities across the pan-proteome panel *T* are transformed to an affinity-proportional scale 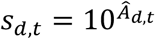 and normalized to obtain a probability-like distribution

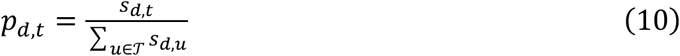

Concentration of this distribution is quantified using the normalized Shannon entropy,

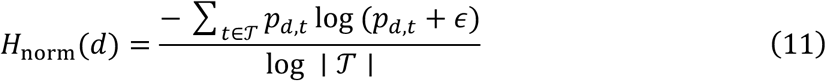

which is converted into an entropy-derived selectivity term 1 − *H*_norm_(*d*), such that larger values indicate stronger concentration of predicted affinity on a limited subset of targets. We further compute a concentration-based measure using the complement of the Gini-Simpson index 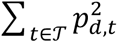, which increases as affinity mass becomes concentrated. Drug potency and separation are summarized using the maximum predicted binding affinity max _*t*_ *Â_d,t_*, the difference between the highest and second-highest predicted affinities, and the ratio between these two values. Each component is scaled across drugs to a common range and combined via a weighted aggregation to yield a composite drug-centric selectivity score. Target-centric selectivity ranks drugs for a given protein by integrating target-specific predicted affinity with the drug’s overall selectivity profile. For each drug-target pair (*d*, *t*), the target-specific affinity is defined as the LinkD-predicted binding affinity *Â_d,t_*. To assess the relative prominence of this interaction within the drug’s proteome-wide binding profile, we compare *Â_d,t_* with the strongest predicted interaction of that drug across the target panel, max 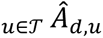, yielding a proteome-relative affinity difference and a corresponding affinity ratio. These quantities indicate whether a target represents the dominant predicted interaction for a drug or is secondary to other predicted targets. The target-centric selectivity score integrates the target-specific affinity, the proteome-relative affinity difference, and the proteome-relative affinity ratio with the drug-level selectivity descriptors derived from the drug-centric formulation, namely, the entropy-derived selectivity and the concentration-based index described above. All components are scaled across drug-target pairs and combined via a weighted aggregation to produce a composite target-centric selectivity score, which is used to rank candidate drugs for each target.

Together, the drug-centric and target-centric formulations provide a unified, proteome-scale quantification of selectivity that jointly captures binding strength, distributional concentration, and competitive separation, enabling systematic prioritization of selective drug-target interactions. To interpret selectivity in a disease-relevant context, targets were annotated as oncogenes, tumor suppressor genes (TSGs), or dual-role genes based on curated cancer gene resources. Selectivity metrics were analyzed within and across these functional classes to assess whether high-selectivity predictions preferentially engage therapeutically actionable targets while minimizing undesirable off-target interactions. Model evaluation emphasized enrichment of known drug-target interactions among high-selectivity predictions, as well as concordance with independent structural, functional, and clinical validation signals.

### Molecular docking simulations

To provide structure-level validation of LinkD predictions, molecular docking was performed for selected drug-target pairs. Protein binding pockets were identified using predicted or experimentally resolved structures. On-target docking scores were compared against off-target docking scores to assess whether LinkD-predicted selectivity is supported by structural binding preferences. The workflow encompasses drug and protein preparation, binding site identification, and docking simulations, integrating several open-source tools to ensure scalability and reproducibility. Detailed process descriptions are provided in the supplementary information.

### Disease target association and target prioritization

We constructed a multi-source cancer target prioritization dataset that integrates disease-target association evidence from the Open Targets Platform^35^, cancer gene role annotations from OncoKB^36^, the COSMIC Cancer Gene Census^37^, and genetic and evolutionary target features from GETdb^38^. The pipeline is designed to enable systematic prioritization of candidate drug targets stratified by cancer type, gene role (oncogene, tumor suppressor gene, or dual role), and druggability. The disease-target association scores were retrieved via the Open Targets GraphQL API for 50 cancer types (ICD-10 C00-C99). For each disease-target pair (*d, t*), the platform computes an overall association score *S*(*d, t*) ∈ [0, 1] by integrating seven evidence sources (genetic associations, somatic mutations, transcriptomics, known drugs, text mining, animal models, and pathway annotations) using a harmonic sum formulation. Pairs with *S*(*d, t*) ≥ 0.05 were retained, a threshold corresponding to the 79th percentile of the score distribution (median S = 0.009, IQR = 0.003-0.037), yielding 9,722 unique targets and 30,797 pairs across 50 diseases. For each target, its oncogenic role was annotated using the OncoKB Cancer Gene List, an FDA-recognized knowledge base that classifies genes as oncogenes, tumor suppressor genes (TSGs), or both, and is cross-validated against the COSMIC Cancer Gene Census and GETdb. Across the 9,722 cancer targets, 479 genes were annotated as oncogenic drivers: 298 oncogenes, 145 TSGs, and 36 with both roles. The remaining 9,243 targets were classified as non-driver. Targets were scored across four domains using the Open Targets target prioritization framework. Precedence was defined by the maximum clinical trial phase achieved. Tractability was assessed by membrane localization, secretion status, ligand availability, small-molecule co-crystallization, and predicted druggable pockets. Doability incorporated mouse ortholog identity (≥80%) and chemical probe availability. Safety integrated genetic constraint (gnomAD LOEUF), mouse knockout phenotype severity, gene essentiality (DepMap), curated adverse events, cancer driver status, paralogue identity, tissue specificity, and tissue distribution, scored on a -1 (unfavorable) to +1 (favorable) scale. A composite score was calculated as the weighted mean of domain-level scores (Precedence 0.20, Tractability 0.30, Doability 0.15, Safety 0.35) and used to rank targets within each cancer type and gene-role stratum.

### Statistical correlation analysis between drug perturbation and CRISPR gene dependency data

Drug-response profiles, quantified as the area under the dose-response curve (AUC), were obtained from large-scale pharmacogenomic screening datasets, including the PRISM Repurposing dataset (PRISM Repurposing Public 24Q2; 1,285 compounds), the Genomics of Drug Sensitivity in Cancer (GDSC; Sanger GDSC1 and GDSC2; 315 compounds), and the Comparative Toxicogenomics Database-annotated PRISM compounds (PRISM_CTD; 336 compounds), totaling 1,936 unique compounds. After quality control and cell-line matching to DepMap identifiers, drug-response measurements spanned 960 GDSC cell lines with valid tissue annotations across 13 cancer tissue lineages. Within each dataset, drug-response values were standardized to z-scores to mitigate platform-specific scale differences.

CRISPR-Cas9-based gene dependency scores were obtained from the Cancer Dependency Map (DepMap, 25Q1 release). Gene fitness effects were estimated using the Chronos algorithm, which models cell population dynamics to account for variable sgRNA efficacy and incomplete knockout penetrance. Gene effect scores were subsequently corrected for copy-number-associated cutting toxicity and screen-quality confounders following the standard DepMap processing pipeline. Chronos gene effect scores were available for 1,178 cancer cell lines across 17,916 genes, with more negative scores indicating stronger dependency.

Drug-response and gene-dependency datasets were harmonized at the cell-line level using unique DepMap identifiers. Only cell lines with both valid drug-response measurements and corresponding Chronos dependency scores for the annotated target gene were retained for downstream analysis. Depending on the specific drug-target pair and data source, the number of matched cell lines ranged from 15 to 605, with a median of 327 per analysis. Missing values in either drug-response or gene-dependency measurements were handled using complete-case analysis. Cell lines lacking valid measurements for a given drug or target gene were excluded from the corresponding correlation analysis but retained for other drug-target evaluations where data were available. No imputation was performed, as missingness primarily reflected incomplete screening coverage rather than systematic measurement failure.

Functional concordance between pharmacological perturbation and genetic dependency was quantified by computing the Pearson correlation coefficient between drug response and the Chronos dependency score of the corresponding target gene across matched cell lines. Correlation analyses were performed independently for each drug-target pair using two-sided tests. To ensure statistical robustness, drug-target pairs evaluated in fewer than 15 matched cell lines were excluded from analysis. Statistical significance was assessed using nominal *P* values derived from the Pearson correlation test. To account for multiple hypothesis testing across all evaluated drug-target pairs, *P*-values were adjusted using the Benjamini-Hochberg procedure to control the false discovery rate (FDR). Drug-target associations with FDR-adjusted *P* < 0.05 were considered statistically significant. High-selectivity drug-target pairs were expected to exhibit stronger concordance between pharmacological sensitivity and genetic dependency across cell lines, consistent with functional target engagement rather than indirect or off-target effects.

### Disease cohort definition using electronic health records

Two electronic health record (EHR) datasets were used: the Mount Sinai Data Warehouse (MSDW) and the UK Biobank (UKB). In the MSDW database, cancer cases were identified using ICD-9-CM and ICD-10-CM codes for malignant neoplasms in an OMOP-formatted EHR database. To improve diagnostic specificity, cancer cases required at least 5 cancer-related diagnosis records or a diagnosis accompanied by cancer-directed treatment. Control participants were drawn from the same population with no recorded cancer diagnosis and at least 1 year of longitudinal follow-up. In the UK Biobank cohort, cancer cases were identified among 501,978 participants using linked primary care, diagnostic, and hospital episode data; controls were defined as participants without a recorded cancer diagnosis prior to the index time. For cancer cases, the index time was defined as the first recorded cancer diagnosis or initiation of cancer-directed therapy, whichever occurred earlier. For controls, the index time was defined as 1 year prior to the last recorded clinical encounter.

### Drug-disease association analysis in EHR

Drug-disease associations were evaluated using electronic health record data from the MSDW and UKB databases. Drug ingredients were harmonized to ChEMBL identifiers, and disease phenotypes were defined using primary ICD-10 diagnosis codes, where “primary” denotes the diagnosis listed as the principal reason for the clinical encounter rather than secondary or historical problem list entries. The resulting analysis set comprised 11.5 million individuals, 1,394 compounds, and 1,783 diseases. For each drug-disease pair, individuals prescribed the test drug prior to disease onset were classified as exposed, while individuals with no recorded exposure formed the unexposed group. Individuals with the disease documented before the first drug exposure were excluded. Drug-disease pairs were required to have >5 disease cases among exposed individuals to ensure model stability. To control for baseline confounding, exposed individuals were matched to unexposed controls using propensity score-based background matching (1:2), with scores estimated from age, sex, and race/ethnicity. Association strength was estimated using logistic regression on the matched cohorts:

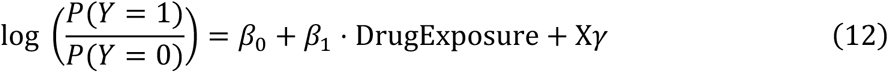

where *Y* denotes disease occurrence and *X* denotes demographic covariates. Odds ratios and 95% confidence intervals were reported.

Multiple testing across all drug-disease pairs was controlled using Benjamini-Hochberg false discovery rate (FDR) correction. For associations that passed FDR thresholds, robustness was further assessed using 10-fold permutation testing, in which exposure labels within matched cohorts were randomly permuted to derive empirical *P*-values. All EHR-based analyses are observational and do not imply causality.

### LinkD AI Agent Architecture and Design

The LinkD agent is an AI system designed to integrate mechanistic, functional, and clinical evidence for systematic analysis of drug-target-disease relationships. It combines a deterministic database layer with a large language model (LLM)-based reasoning and planning module, enabling natural language querying and autonomous multi-step analysis. The underlying knowledge base integrates 276,147 drug-target-disease associations from clinical trial records (4,274 drugs, 1,520 targets, and 2,684 diseases), 13,008 curated causal gene-disease relationships (covering 3,400 genes and 3,859 diseases), and functional annotations for 1,029 oncogenes and tumor suppressor genes (485 oncogenes, 404 tumor suppressors, and 140 with dual roles).

Mechanistic evidence is provided through large-scale predicted drug-protein binding affinity and selectivity profiles for 14,981 drugs across 20,385 human protein targets, represented as *pK*_*d*_values (range 2.90-10.95) and three selectivity categories (Highly Selective, Moderate poly-target, and Broad-spectrum). Functional relevance is assessed using correlations between CRISPR/Cas9 gene dependency scores and drug response metrics (AUC score) from GDSC, PRISM, and PRISM_CTD datasets, comprising 464,820 correlation records across 1,109 drugs and 17,029 genes. Real-world clinical evidence is incorporated from electronic health record cohorts, including the Mount Sinai Health System (41,120 drug-disease associations across 730 drugs and 964 ICD-10 codes, summarized using logistic regression odds ratios) and the UK Biobank (693 drug-disease associations across 143 drugs and 20 ICD-10 codes, summarized using odds ratios), each under predefined statistical thresholds.

User queries are parsed by the LLM agent to extract biomedical entities (drug IDs, gene symbols, disease names, ICD codes, and clinical trial phases) and intent, and complex requests are decomposed by a planning agent into sequential analytical steps. Each step executes structured database queries with explicit quantitative criteria, and results are aggregated across modalities into unified summaries. All numerical statistics are derived directly from structured data sources, while the LLM is used solely for query classification, plan generation, and result interpretation. The agent is deployed through a Gradio-based interactive web interface that supports transparent, reproducible, and scalable evidence integration.

### Cell viability assay

Human prostate cancer LNCaP cells were obtained from ATCC (Manassas, VA). Cells were cultured in RPMI1640 supplemented with 10% FBS, 2 mM L-glutamine, and 1x antibiotic/antimycotic (Gemini Bio-Products, Sacramento, CA) at 37° C in 5% CO2. Cells were authenticated by human short tandem repeat profiling and Mycoplasma testing was performed using the MycoAlert PLUS Assay. Cell viability was assessed using the MTT assay (3-[4,5-dimethylthiazol-2-yl]-2,5-diphenyl tetrazolium bromide; Invitrogen) as previously described^39^. Briefly, LNCaP cells were seeded at a density of 2.5 × 10³ cells per well in 96-well plates containing RPMI medium supplemented with 10% fetal bovine serum (FBS). Cells were treated with either DMSO (vehicle control), enzalutamide (10 µM), propranolol (100 µM), carvedilol (50 µM), or combinations of enzalutamide with propranolol or carvedilol for three days. The concentration of propranolol and carvedilol were used based on the prior literature^40,41^. Cell growth experiments were performed in biological duplicates, with eight wells per condition in each experiment. After three days of treatment, cells were incubated with MTT (0.5 mg/mL) for 1 h at 37° C in 5% CO2. Media was removed and formazan crystals were then dissolved in 100 μL/well isopropanol for 10 minutes, and absorbance was measured at 570 nm using a BioTek plate reader. Blank cell free wells with isopropanol were used as a background control.

## Data availability

All data used in this study were obtained from publicly available resources or generated as part of this work. Clinical trial-derived drug-target-disease associations, causal gene-disease relationships, and oncogene annotations were obtained from publicly accessible databases, as described in the Methods. Drug-protein binding affinity and selectivity predictions generated by the LinkD framework, as well as processed drug response and electronic health record (EHR) summary statistics, are available at Zenodo website (https://zenodo.org/records/19241152). Access to individual-level EHR data is restricted due to patient privacy and data-use agreements; only aggregated, de-identified statistical results were used in this study.

## Code availability

The source code and data for the LinkD-Agent, including database integration modules, agent planning and reasoning components, and the interactive web interface, is available at https://github.com/mmetalab/LinkD and https://linkd-agent.onrender.com. The repository includes documentation, configuration files, and example queries needed to reproduce the analyses reported in this study. Scripts for molecular docking, binding affinity integration, and agent execution are provided. Model weights and API keys for third-party large language models are not distributed and must be obtained separately in accordance with the providers’ terms of use.

## Acknowledgements

This research was partially funded by the National Institute of Health (R01CA274967 to G. Chakraborty). G Chakraborty was supported by a Prostate Cancer Foundation Young Investigator Award. The authors thank all members of the Huang lab for constructive discussion. Large language models (LLMs) may have been used in the initial drafts of coding and writing of this work. All final codes and texts have been extensively edited and verified by the authors. This work was supported in part the Mount Sinai Data Warehouse (MSDW) resources and staff expertise provided by Scientific Computing and Data at the Icahn School of Medicine at Mount Sinai and supported by the Clinical and Translational Science Awards (CTSA) grant UL1TR004419 from the National Center for Advancing Translational Sciences. Research reported in this publication was also supported by the Office of Research Infrastructure of the National Institutes of Health under award number S10OD026880 and S10OD030463. The content is solely the responsibility of the authors and does not necessarily represent the official views of the National Institutes of Health. This work was supported by NIH NIGMS R35GM138113, NIH NIGMS 2R35GM138113, NIA UG3AG105083, and ACS RSG-22-115-01-DMC to K.H.

